# Geographic and subsequent biotic isolations led to a diversity anomaly of *Heterotropa* (Aristolochiaceae) in insular versus continental regions of the Sino-Japanese Floristic Region

**DOI:** 10.1101/2020.04.25.060632

**Authors:** Daiki Takahashi, Yu Feng, Shota Sakaguchi, Yuji Isagi, Ying-Xiong Qiu, Pan Li, Rui-Sen Lu, Chang-Tse Lu, Shih-Wen Chung, Yang-Shan Lin, Yun-Chao Chen, Atsushi J. Nagano, Lina Kawaguchi, Hiroaki Setoguchi

## Abstract

The Sino-Japanese Floristic Region is highly diverse with respect to temperate plants. However, the reasons for this diversity are poorly understood because most studies have only considered geographic isolation caused by climatic oscillations. *Heterotropa* (genus *Asarum*; Aristolochiaceae) diverges here and shows high species diversity in insular systems (63 species) compared to continental areas (25 species). *Heterotropa* shows low dispersal ability with small distribution ranges, implying diversification by geographic events, and high floral diversity, implying pollinator-mediated diversification. To reveal how abiotic and biotic factors have shaped the diversity anomaly of *Heterotropa*, we conducted phylogenetic analysis using ddRAD-seq and chloroplast genome datasets including 79 species, estimation of floral trait evolution, and comparison of isolation factors within clades based on distribution range and floral trait analysis. Phylogenetic analysis indicates that *Heterotropa* originated in mainland China and expanded to the Japanese Archipelago in the Miocene, and the major clades almost correspond to geographic distributions. Floral traits evolved repeatedly in the tip nodes within the clades. Although the major clades include a high proportion of species pairs showing isolation by floral traits, there are no conditional relationships between two isolation factors, indicating that most species pairs with floral trait isolation are distributed allopatrically. The repeated exposure and submergence of land-bridges caused by climatic oscillations would have led to significant population fragmentation in insular systems. Thus, the diversity anomaly of *Heterotropa* would have resulted from geographic and climatic events during the Miocene, while subsequent repeated floral trait evolution would have followed geographic isolation during the Pleistocene.

## Introduction

The Sino-Japanese Floristic Region (SJFR) extends from the eastern Himalayas to the Japanese Archipelago through south and central China (Takhtajan, 1969; Wu & Wu, 1996). This region can boast one of the most diverse temperate floras anywhere in the world and has high endemism (Wu & Wu, 1996). This diversity has been thought to be linked to climatic and physiographical complexity and historical environmental changes associated with the Pleistocene (< 2.6 Mya) climatic oscillations (Qian & Ricklefs, 2000). During glacial periods, when the climate of this region was cooler by ca. 4-6 °C and sea level was approximately 130m lower than its present level, temperate plants in this region retreated to refugia at lower altitudes or southern parts (Harrison, Yu, Takahara, & Prentice, 2001; Tsukada, 1984). During the interglacial periods, expansion to higher altitudes or northern parts would have occurred. In the eastern island systems, sea level changes due to climatic oscillations have caused repeated formation and division of land-bridges in the East China Sea (Ujiie, 1990), and these events have provided opportunities for population expansion and fragmentation (Qiu, Fu, & Comes, 2011). Many phylogeographic studies have revealed that the present interspecific and intraspecific genetic structures of temperate plants in this region reflected the range shifts caused by climatic oscillations (Li, Yan, & Ge, 2012; Sakaguchi et al., 2012; Setoguchi et al., 2006; Wang et al., 2015; Yang et al., 2017). It has been considered that these climatic and associated environmental changes during the Pleistocene triggered range fragmentation, vicariance, and population isolation (Qiu et al., 2011). It has also been hypothesised that allopatric speciation would be a major mode of speciation in the temperate plants of this region (Qian & Ricklefs, 2000).

Although the importance of geographic isolation as a major isolation mechanism in plants has been addressed (Boucher, Zimmermann, & Conti, 2016; Esselstyn, Timm, & Brown, 2009; Govindarajulu, Hughes, & Bailey, 2011; Verboom, Bergh, Haiden, Hoffmann, & Britton, 2015), recent studies in other regions have implied that biotic factors also promote species diversification (Lagomarsino, Condamine, Antonelli, Mulch, & Davis, 2016; L. Lu et al., 2019). In particular, floral trait evolution has been thought to promote speciation through segregation of gene flow by pollinator shifts (Armbruster, 2014; Van der Niet, Peakall, & Johnson, 2014). Previous studies have shown that the tempo and pattern of floral trait evolution varies distinctively among lineages, and floral trait evolution has been implicated in shaping patterns of species diversification (Givnish et al., 2015; Jaramillo & Manos, 2001). It has been considered that biotic drivers can play complementary roles to abiotic factors in reproductive isolation; that is, biotic factors can facilitate reproductive isolation even without geographical isolation (Rundle & Nosil, 2005). To fully understand the diversification process of species groups, it is essential to reveal the relative contributions of biotic and abiotic drivers. However, many studies conducted in the SJFR have only discussed the role of allopatric fragmentation due to geographic and climatic events, and few studies have considered other factors as drivers of the diversification of the temperate plants. In addition, although the high endemism in the SJFR provides an outstanding opportunity for investigating the evolutionary history of plant diversification (Yang et al., 2017), to date, most phylogeographic studies in the region have focused on individual species or only small groups, including fewer than 10 taxa (But Mitsui et al., 2008; Yoichi, Jin, Peng, Tamaki, & Tomaru, 2017). Thus, our knowledge of the diversification process of temperate plants in the SJFR remains fragmentary, due to a lack of integrative multidimensional studies of morphology, phylogeny, biogeography, and ecology with adequate sampling of diversified groups.

In this study, we focused on the *Heterotropa* (genus *Asarum*; Aristolochiaceae) as a model group endemic to the SJFR, and one of the most speciose warm-temperate plant groups (comprising approximately 90 species) in this region (Sugawara, 2006). Taxa of *Heterotropa* are low-growing, rhizomatous herbs that grow in shaded understories, and are distributed in mainland China (25 species: Huang, Kelly, & Gilbert, 2003), Taiwan (13 species: C. T. Lu & Wang, 2009; Chang Tse Lu & Wang, 2014), and the Japanese archipelago, including the Ryukyu islands (52 species: Sugawara, 2006). The species diversity of *Heterotropa* is uneven, and given the difference in areas, *Heterotropa* shows higher diversity in the eastern insular region (from Taiwan to mainland Japan; 2.7 × 10^−4^ species/km^2^ and mainland China; 1.3 × 10^−5^ species/km^2^, see Results for details). Some taxa of *Heterotropa* have very limited geographic ranges (e.g., in only one island or mountain range), and the dispersal ability of *Heterotropa* is estimated to be 10□50 cm per year due to its myrmecochore seeds with elaiosome (Hiura, 1978; F. Maekawa, 1953). Low dispersal ability promotes genetic differentiation among populations and often leads to allopatric speciation (Petit et al., 2005). These confined distribution ranges and the low dispersal ability led us to hypothesise an allopatric speciation process for *Heterotropa*. On the other hand, *Heterotropa* taxa are characterised by high divergence in floral traits, in terms of their shapes, sizes, and colours of calyx tubes and lobes (Fig. 1a & S1), while their vegetative traits show almost no differences (Sugawara & Ogisu, 1992). The sepals connect beyond attachment to the ovary and form a calyx tube with calyx lobes (Sugawara, 1987), and their flowers have been hypothesised to mimic fungi in order to attract fungus gnats (Sinn, Kelly, & Freudenstein, 2015; Vogel, 1978). In addition to flower shape, *Heterotropa* taxa are highly divergent in flowering time; most taxa have flowers in spring, while others have flowers in autumn or winter (Huang et al., 2003; Sugawara, 2006). A genus-wide phylogenetic study of *Asarum* showed that diversification of *Heterotropa* could have been triggered by the presence of putative fungal-mimicking floral structures, loss of autonomous selfing, and loss of vegetative growth (Sinn et al., 2015). Given these characteristics, we considered that *Heterotropa* would be an ideal subject for investigating the relative importance of the abiotic and biotic effects on its diversification in the SJFR. Our previous phylogenetic study using the ITS region showed that *Heterotropa* was monophyletic and comprised two clades, which corresponded to species distribution ranges, namely mainland China and the island arc from Taiwan to mainland Japan, and the estimated time of divergence between these two clades was approximately 9 million years ago (Mya) in the late Miocene (Takahashi & Setoguchi, 2018). In order to obtain the taxonomic implications of insular *Heterotropa*, Okuyama et al., (in press) conducted phylogenetic analysis using RAD-seq datasets including 47 insular and 5 continental species, and they implied that insular *Heterotropa* comprises nine groups that almost correspond to the geographic distributions. However, due to the low resolution of the datasets or lack of inclusive sampling around the SJFR, the formation mechanisms of the diversity anomaly of *Heterotropa* in insular systems and their diversification history in terms of temporal and spatial patterns of floral trait evolution (e.g., parallel evolution in each region or single origin with subsequent expansion) remain unknown.

**Fig. 1.**
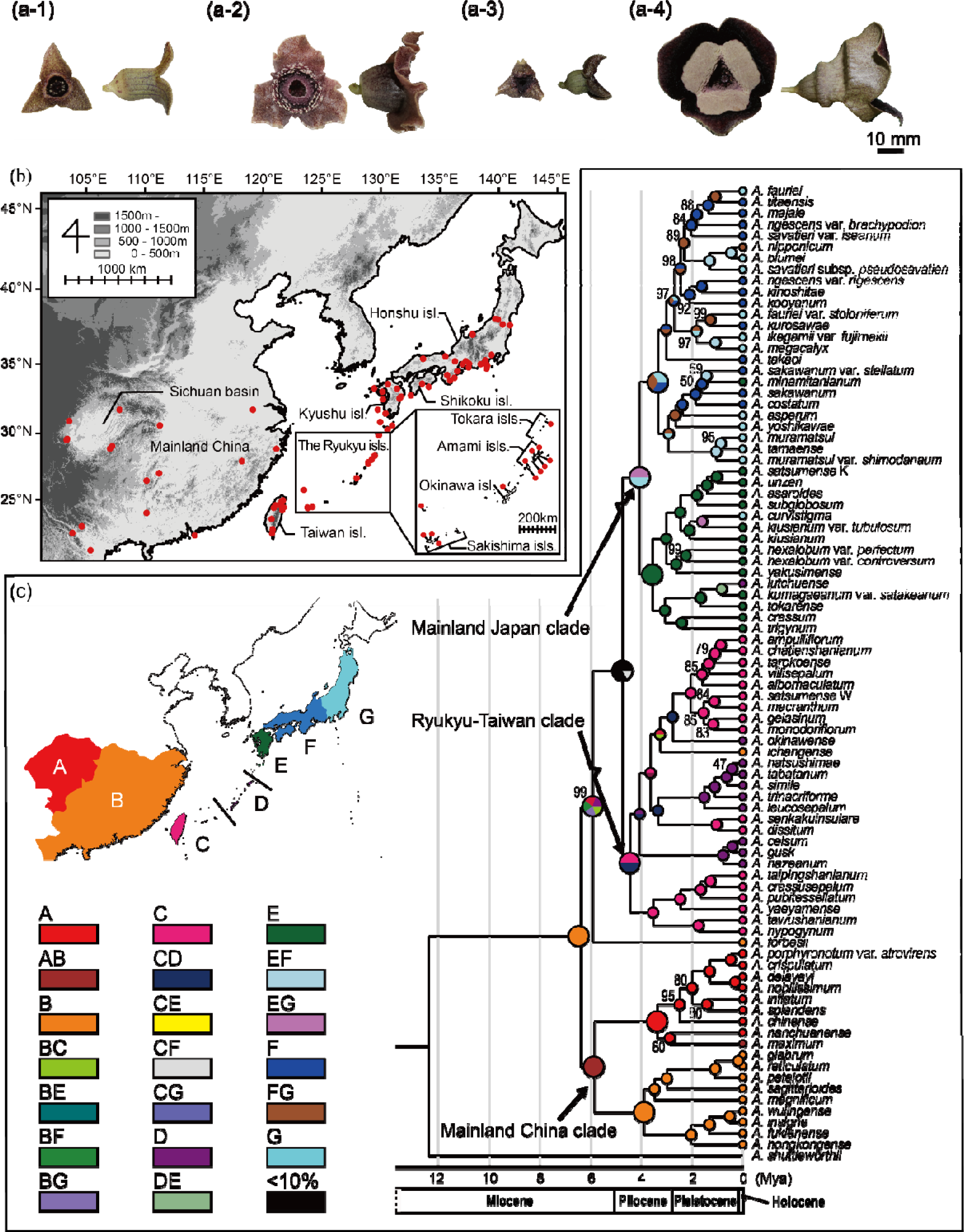
The diversification of *Heterotropa* in the Sino-Japanese Floristic region. (a) Photographs of the flowers of *Heterotropa* taxa (a-1; *Asarum takaoi*, a-2; *Asarum unzen*, a-3; *Asarum dissitum*, and a-4; *Asarum maximum*). (b) Map of the Sino-Japanese Floristic Region. Red circles indicate sampling points. (c) Time-calibrated molecular phylogenetic tree and the ancestral areas estimated from Bayesian analysis and statistical dispersal-vicariance analysis (S-DIVA). Posterior probabilities of < 100% are indicated above or below branches, and the branches without posterior probabilities are those with 100% support. The colours of pie charts reflect the estimated distribution areas according to the biogeographic delimitation as in map (A; Sichuan basin and surrounding mountains, B; other parts of mainland China, C; Taiwan and southern Ryukyu islands, D; central Ryukyu islands, E; northern Ryukyu islands and Kyushu island, F; southern part of mainland Japan including Shikoku island, and G; northern part of mainland Japan).

Our primary purpose in this study was to reveal the diversification history of *Heterotropa* taxa in the SJFR, especially focusing on the relative contribution of abiotic and biotic drivers and diversity anomalies in insular systems. For this purpose, we first constructed a highly resolved and time-calibrated phylogenetic tree including most *Heterotropa* taxa using genome-wide single nucleotide polymorphisms (SNPs) and chloroplast genomes. We also estimated the patterns of floral trait evolution using macroevolutionary modelling and indices of phylogenetic signals based on the obtained phylogenetic tree. In addition, to test whether the geographic isolations would be major forces for speciation or whether biotic factors were more likely to facilitate the speciation in *Heterotropa*, we compared patterns of geographic and morphological isolation within the clades and between sister taxa pairs and non-sister taxa pairs. The implications for the relative roles of biotic and abiotic drivers and the temporal and spatial patterns of their influences would help to understand the diversification process of temperate plants in the SJFR.

## Materials and methods

### Taxon sampling

Here, we present a comprehensive sample collection of sect. *Heterotropa*, increasing the number of species from 64 species (Takahashi & Setoguchi, 2018) or 52 species (Okuyama et al., in press) to 79 species (Fig 1b, Table S1), including three undescribed but morphologically distinct taxa (*Asarum kiusianum* var. *tubulosum* nom. nud. [Maekawa, 1983], *Asarum tarokoense* nom. nud. [Lu et al., in prep.], and *Asarum titaense* nom. nud. [Setoguchi et al., in prep]) in our analysis. *Asarum satsumense* has been considered to be distributed in both Taiwan and Kyushu islands (C.-T. Lu, Chiou, Liu, & Wang, 2010), and our preliminary examination suggested that the Taiwanese entity should be distinguished from the species (Lu and Takahashi, personal observation). Thus, in this study, we treated *A. satsumense* collected from the Taiwan and Kyushu islands as a different taxon (*A. satsumense* W; Taiwanese and *A. satsumense* K; Japanese). The sample set included 46 Japanese species (52 taxa), 13 Taiwanese species (13 taxa), and 20 mainland Chinese species (20 taxa). As an outgroup species, we used one sect. *Hexastylis* species (*Asarum shuttleworthii*) according to our previous study (Takahashi & Setoguchi, 2018).

### Chloroplast genome sequencing and phylogenetic analysis

For chloroplast genome construction, we conducted genome sequences of five *Asarum* species, including *A. shuttleworthii* (Tables S2 & S3). Details of library preparation, sequencing methods, and phylogenetic analysis are described in Appendix 1.

### Double-digest restriction-associated DNA sequencing

Genomic DNA was extracted from silica-dried leaf tissues using the CTAB method (Doyle & Doyle, 1987). For all collected samples, a double-digest restriction-associated DNA (ddRAD) library was prepared using Peterson’s protocol with slight modifications (Peterson, Weber, Kay, Fisher, & Hoekstra, 2012). Genomic DNA was digested with BglII and EcoRI, ligated with Y-shaped adaptors, amplified by PCR with KAPA HiFi HS ReadyMix (KAPA BIOSYSTEMS) and size-selected with the E-Gel size select (Life Technologies, CA, USA). Approximately 350 bp of library fragments were retrieved. Further details of the library preparation method were described in a previous study (Sakaguchi et al., 2015). Sequencing was performed with paired-read 101bp + 100bp mode of HiSeq2500 (Illumina, CA, USA).

### Data analysis

The raw reads obtained from ddRAD-seq were trimmed by Trimomomatic v. 0.32 software (Bolger, Lohse, & Usadel, 2014) with the following settings: HEADCRAP:10, LEADING:30, TRAILING:30, SLIDINGWINDOW:4:30, AVGQUAL:30, and MINLEN:50. The program ipyrad (http://github.com/dereneaton/ipyrad) was used to process the ddRAD-seq reads and detect SNPs. The parameters that influenced the assembly were set as follows: the minimum depth coverage for base calling at each locus was set at 6 and the similarity threshold for clustering reads within/across samples was set at 0.85. Potential paralogous loci were filtered out based on the number of samples with shared heterozygous sites (more than 15 sites). We explored a range of thresholds for the minimum genotyped samples (30, 51, and 70 samples; equivalent to 50%, 75%, and 90% of samples were genotyped, respectively). All three data sets were examined in the phylogenetic analysis, and we selected the 50% genotyped data set as the primary data set for all other analyses (see Results section).

### Phylogeny construction and dating

To construct a phylogenetic tree of ddRAD-seq data and estimate divergence times within the *Heterotropa* clade, we used Bayesian inference (BI) in BEAST v. 1.10.4 (Drummond & Rambaut, 2007). We calibrated the crown age of the *Heterotropa* clade (with lower limit of 4.77 Mya and upper limit of 14.54 Mya) following the results of the chloroplast genome phylogenetic analysis (see Results section). The Markov Chain Monte Carlo method was performed using two simultaneous independent runs with four chains each (one cold and three heated), saving one tree every 1000 generations for a total 30,000,000 generations with 10% burn-in for each run. The convergence of the chains was checked using the program Tracer v. 1.5 (Rambaut & Drummond, 2013).

### Ancestral area reconstruction

To infer the ancestral areas and phylogeographic history of *Heterotropa* taxa, we performed statistical dispersal-vicariance analysis (S-DIVA) using RASP v3.2 (Yu, Harris, & He, 2010). The analysis was conducted using a phylogenetic tree of the ddRAD-seq dataset with a default setting. We divided the distribution range of *Heterotropa* into seven regions: (A) Sichuan basin and surrounding mountains, (B) other parts of mainland China, (C) Taiwan and southern Ryukyu islands, (D) central Ryukyu islands, (E) northern Ryukyu islands and Kyushu island, (F) southern part of mainland Japan including Shikoku island, and (G) northern part of mainland Japan (see Fig. 1c). The boundaries of these regions were defined with reference to biogeographic studies (e.g., C and D; Kerama gap, D and E; Tokara gap [Kimura, 1996], F and G; Itoigawa-Shizuoka tectonic line [Okamura *et al*., 2017]) and phylogeographic studies (Landrein, Buerki, Wang, & Clarkson, 2017).

### Analysis of trait evolution

To estimate the evolutionary patterns of floral traits concerning reproductive barriers, we focused on flowering time, which we defined as a month when the taxon starts flowering, and calyx tube width, defined as a median value between maximum and minimum calyx tube widths. Flowering time is related to the local environment, including pollinator fauna, and its differences play a role in the reproductive barrier among taxa. Calyx tube width would be linked to pollinator size selection and its difference could affect the difference in pollinator fauna, which leads to reproductive isolation. The trait values were obtained from field observations and literature (Huang et al., 2003; C. T. Lu & Wang, 2009; Sugawara, 2006). The evaluated trait values are summarised in Table S1.

We first plotted each trait value on the tips of the phylogenetic tree to visualise trait-phylogeny relationships using “plotTree.wBars” function in “phytools” package (Revell, 2012) for R v. 3.5.4 (R Core Team, 2013). Next, to estimate the tempo and mode of floral trait evolution on the phylogenetic tree of *Heterotropa*, we conducted Bayesian macroevolutionary analysis for flowering time and calyx tube width implemented in BAMM v. 2.5.2 (Rabosky, 2014). BAMM models shift in macroevolutionary regimes across a phylogenetic tree using reversible-jump Markov chain Monte Carlo (rjMCMC) sampling. To treat the flowering times as continuous traits, we transformed the flowering times of *Heterotropa* onto a circular scale, where the difference between December and January is equivalent to one month. The prior values were set using the package “BAMMtools” (Rabosky et al., 2014) for R. To ease model complexity, we adopted the time-invariant Brownian motion model of trait evolution. The analysis involved a rjMCMC run of 10,000,000 generations sampled every 10,000 steps, and the initial 3,000,000 generations were discarded as burn-in. The rjMCMC convergence was confirmed using BAMMtools. For each trait, we reported the mean scaled tree from the outputs of BAMM, in which each branch length is shortened or stretched proportional to the mean evolutionary rates of the trait using the “getMeanBranchLengthTree” function in BAMMtools. In addition, to infer the temporal change in traits’ evolution rates, we plotted the evolutionary rate variation through time using “plotRateThroughTime” function for each trait and clade.

To determine the evolutionary pattern of the two floral traits within *Heterotropa* taxa, we also measured phylogenetic signals by estimating κ and λ Pagel’s statistics (Pagel, 1999). These values provide insights into temporal patterns of trait evolution. Values of κ and λ close to 1 indicate that the trait showed a strong phylogenetic signal and evolved in a Brownian motion-like manner. Values κ and λ close to 0 indicate that closely related taxa showed relatively row trait similarity, implying that the trait evolved punctually and that the trait changes occurred late in evolutionary history. Estimation of Pagel’s statistics was conducted by using the “phylosig” function in the “phytools” package. We tested for the significance of phylogenetic signal values using 999 randomisation tests.

### Geographic overlaps within each clade

The potential geographic range of each taxon was set by creating a convex polygon from the distribution data. Distribution data for all taxa were based on the specimen records of the herbarium of Kyoto University (KYO), S-Net data portal (http://science-net.kahaku.go.jp/, accessed on 2019/1/26), and the Chinese Virtual Herbarium (http://www.cvh.org.cn/, accessed on 2019/1/26), and the results of personal observations. Because the distribution range of many *Heterotropa* taxa tends to be confined to a small area, we did not set any threshold values of the number of records to make distribution data. In total, we collected 1396 occurrences, with a mean number of 16 records per taxon (minimum 1, maximum 197). We were not able to find available records for *Asarum nobillisimum* and excluded this taxon from the analysis. A single convex polygon for each taxon was created by connecting the outline of the occurrence point(s) by placing a 1 km round buffer and masking by a costal line. Following Anacker and Strauss (2014), we calculated range overlap as the area occupied by both taxa divided by that of the smaller ranged taxa. The range overlap values ranged from 0 (no overlap) to 1 (complete overlap) and were calculated for all pairs of taxa within the three major clades (mainland China, Ryukyu-Taiwan, and mainland Japan clades, see Results section). All geographic analysis were conducted by using “sf” package (Pebesma, 2018) in R.

### Geographic and morphological isolation

To infer the patterns of reproductive isolation in *Heterotropa*, we measured the geographic and morphologic isolations between OTU pairs. The geographic isolation index was calculated by transforming the geographic overlap values into binary data (0; overlap, 1; non-overlap). We set the values of the geographic isolation index of partially overlapping pairs to 0. We also calculated the morphological isolation indices for two floral traits (flowering time and calyx tube width) between OTU pairs. To estimate the differentiation of the floral traits, we set the intervals of the traits using the first and last flowering months, and the maximum and minimum values of calyx tube width, respectively, according to the trait data. For each trait, when a pair of taxa had an overlap of the intervals, they were scored as 0 for “overlapping”; if they showed no overlaps, they were scored as 1 for “isolation”. We calculated these isolation indices for all pairs of OTUs and conditional relationships among the attributes within the three major clades.

### Comparison of sister and non-sister taxon pairs

If allopatric speciation was common in *Heterotropa*, we can expect that closest relatives would exhibit less geographic overlap compared with non-sister pairs. Likewise, we can speculate whether the floral traits are concerned with speciation by examining how floral traits differentiate between close relatives. We compared the geographic and morphological isolation values of sister taxa with those of non-sister taxa by using Bayesian generalised linear mixed models (GLMM) analysis. GLMM analyses were conducted using “brms” package (Burkner, 2017) in R. We set each isolation index as a response variable following the Bernoulli distribution, and the relationship of pair (sister or non-sister taxa) was used as the fixed effect. We selected sister pairs with posterior probabilities of nodes higher than 80% in the phylogenetic analysis of ddRAD-seq data sets as sister taxa and excluded pairs with less than 80% support in the analysis. We included a nested random effect: OTU pairs nested within their clades (Mainland Japan, Ryukyu-Taiwan, or mainland China). For OTU pairs, to account for non-independence resulted from the use of pairwise matrices of individual-level metrics, we used “multi-membership” function implemented in brms. We run the four chains for 10,000 iterations with a warm-up length 5,000 of iterations with default priors. We also conducted the same analyses using the data set adopting the sister taxa with posterior probabilities > 99% to account for phylogenetic uncertainty in our analyses.

## Results

### ddRAD-seq data

The ddRAD-seq reads generated by Illumina sequencing were deposited in GenBank (BioProject ID: PRJDB8943). After filtering low-quality reads and bases, the number of reads of each sample ranged from 510,135 to 2,558,859 reads and the average number of reads was 1,348,526 (Table S1). Our 50% genotyped matrix consisted of 469 loci, which contained 3,415 parsimony-informative SNPs. Our 70% and 90% matrices included 117 and 46 loci with 710 and 266 parsimony informative SNPs, respectively.

### Phylogenetic inference and ancestral area reconstruction

Phylogenetic analysis based on 59 chloroplast CDS regions supported all clades within the tree with posterior probabilities > 0.99 and showed that the *Heterotropa* was monophyletic (Fig. S2). The estimated divergence time between a mainland China taxon (*A. wulingense*) and the insular clade including *Asarum forbesii* was 9.16 Mya (95% HPD: 4.66 – 13.55 Mya).

Phylogenetic analysis with 50% genotyped ddRAD-seq matrix yielded strongly supported clades within *Heterotropa* (Fig. 1c). Within the *Heterotropa* clade, the mainland China clade diverged early, followed by a splitting off of *A. forbesii*, which is also distributed in mainland China, and the clade that consisted of insular taxa with only one mainland Chinese taxa (*Asarum ichangense*). Within each insular and mainland Chinese clade, there were subclades that almost corresponded with the geographic structures. The mainland China clade was divided into two subclades, including the taxa distributed around the Sichuan basin and in other parts of mainland China. The insular clade consisted of two subclades, including taxa distributed in the southern parts of the Japanese island arc (from Taiwan to the Amami islands; Ryukyu-Taiwan clade) and northern parts (from Tokara islands to Honshu; mainland Japan clade). *Asarum ichangense* was included in the Ryukyu-Taiwan clade. Within the mainland Japan clade, almost all taxa found on Kyushu island (except for *Asarum minamitanianum* and *Asarum asperum*) formed a clade with *Asarum lutchuense* and *Asarum curvistigma*, which are found on Amami islands and Honshu island, respectively. The other taxa in mainland Japan clade split into two subclades, both of which included southern and northern Japanese taxa. All clades mentioned above showed high support (posterior probabilities > 99%) and diverged during pre-Pleistocene periods (> 2.6 Mya; Fig. S3). The topologies within the insular clade of ddRAD-seq data (this study) and ITS (Takahashi & Setoguchi, 2018) were different from those of the chloroplast data. This discordance could be due to incomplete lineage sorting within the recently divergent clades and low mutation rates of chloroplast CDS regions (Wolfe, Li, & Sharp, 1987).

Our BI trees inferred from 75% and 90% genotyped ddRAD-seq matrices also supported the hypothesis that *Heterotropa* split into the insular and mainland China clades, while the nested structures were not resolved (Fig. S4). The tree obtained from the 50% genotyped matrix showed relatively high support of nodes (mean posterior probabilities = 91.0%), while other matrices showed lower supports (75% genotyped matrix; 77.7%, and 90% genotyped matrix; 63.2%). Thus, we adopted the 50% genotyped tree for further analysis.

The results of S-DIVA analysis (Fig. 1c) showed that the origin of *Heterotropa* was in the mainland China region (B) and that dispersal to insular systems occurred subsequently. From insular systems, only one back-dispersal to mainland China was estimated (*A. ichangense*). The common ancestral area of the Ryukyu-Taiwan clade is presumed to be Taiwan and the southern Ryukyu islands (C) or Taiwan with southern and central Ryukyu islands (CD). In the mainland Japan clade, several taxa colonised the northern part of Japan (G) from the southern part (E and F).

### Analysis of trait evolution

Most *Heterotropa* taxa (55 taxa) analysed in this study start flowering in spring (March to June), whereas 11 taxa flower in autumn (September to November) and 19 taxa in winter (December to February). These autumn-flowering and winter-flowering taxa were scattered across all three major clades, and many tip nodes did not share the flowering time (Fig. 2a), implying that the flowering time of *Heterotropa* changed several times in each of the three major clades, especially in tip nodes or branches. With respect to the calyx tube width (Fig. 2b), the taxa that have more than 15 mm of calyx width represented two clades (mainland Japan and mainland China clades), and the changes would have evolved in parallel.

**Fig. 2.**
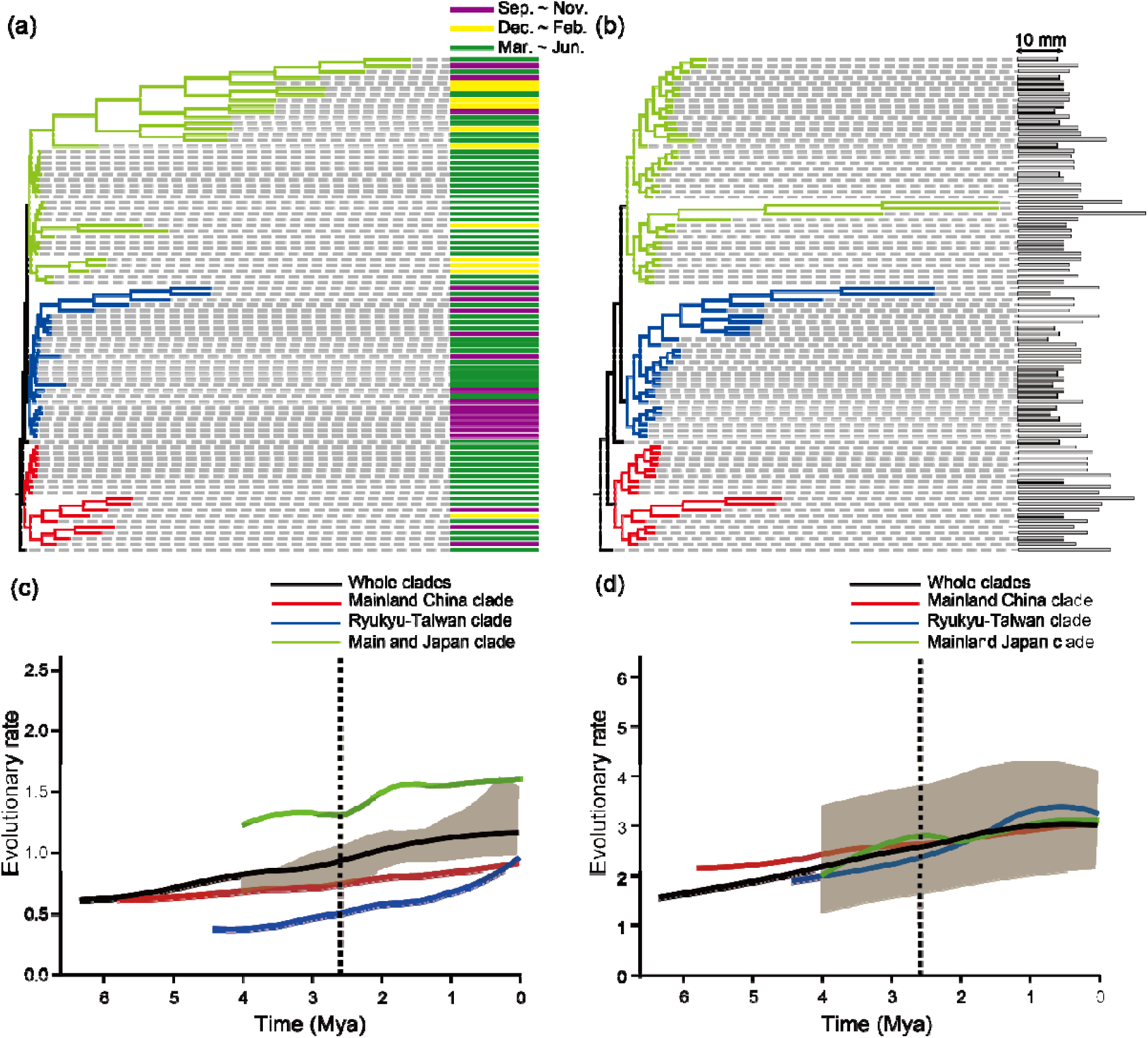
The results of macroevolutionary modelling analysis of floral trait evolution estimated from BAMM. Phylogenetic trees with branch lengths were proportional to their marginal phenotypic evolution rate of flowering time (a) and calyx tube width (b). The topology of the trees is the same as in Figure 2. Colours and lengths of bars across the tips of the phylogenetic trees represent flowering time and mean calyx tube width (mm), respectively. Flowering times are classified into three types: autumn (flowering at September to November; purple), winter (flowering at December to February; yellow), and spring (flowering at March to June; green). The median value with 95% CI of evolutionary rate of flowering time (c) and calyx tube width (d) through time in each clade (black; whole clades, red; mainland China clade, blue; Ryukyu-Taiwan clade, and light green; mainland Japan clade). Dashed vertical lines indicate the boundary between the Pliocene and the Pleistocene (2.6 Mya).

The results of BAMM analysis showed that for both traits, a single macroevolutionary rate was unlikely to fit our genetic data (Fig. S5), indicating that several rate shifts of trait evolution would have occurred in *Heterotropa*. From the ddRAD-seq trees with branch lengths drawn proportional to their marginal evolutionary rates (Fig. 2ab), the rate shifts were distributed across all three major clades within *Heterotropa* for both traits. Most of these shifts accelerated the evolutionary rates, and the shifts were estimated at more terminal nodes containing only several taxa, rather than an early burst of evolutionary rate shift in crown nodes. As an exception, the clade containing half the mainland Japanese taxa showed a substantial increase in the rate of flowering time evolution. Rate variation through time plots indicated that the evolutionary rates of both traits increased in all clades, and sudden increases were observed in the flowering time of the mainland Japan and Ryukyu-Taiwan clades and in the calyx tube width of the Ryukyu-Taiwan clade during the Pleistocene (Fig. 2cd).

For both traits, κ and λ values were relatively low (κ for flowering time; 0.389, for calyx tube width; 0.390 and λ for flowering time; 0.433, calyx tube width; 0.521), while all values were significantly different from 0 (*p* < 0.05), except for λ at flowering time (*p* = 0.311; Table S4). This indicates that for both traits, trait evolution would have occurred later within the regional clades and punctually in the evolutionary history.

### Geographic and floral trait isolation

The mean values of the distribution areas of the taxa within mainland China, Ryukyu-Taiwan, and mainland Japan clades were 172,168 km^2^, 570 km^2^, and 6,524 km^2^, respectively, (Table S1; Fig S6). Within these clades, most of the taxa pairs were distributed allopatrically (Table 1; Fig. S7abc). The mainland China clade contained a relatively low proportion of geographically isolated pairs (69.85%), followed by the Ryukyu-Taiwan clade (88.60%), and mainland Japan clade (90.36%). Within the mainland Japan clade, 52.44% of taxon pairs showed flowering time isolation, whereas the proportion of taxon pairs showing isolation in calyx tube width was lower (36.28%). Conversely, both Ryukyu-Taiwan and mainland China clades contained relatively high proportions of taxa pairs showing calyx tube width isolation (56.13% and 68.63%, respectively), while only 21.37% and 28.76% of taxa pairs showed flowering time isolation, respectively. All three clades included a high proportion of taxa pairs that differentiated in either flowering time or calyx tube width (mainland Japan: 71.02%, Ryukyu-Taiwan: 66.76%, and mainland China: 76.47%). In all three clades, most pairs showing floral trait differentiation were geographically isolated, that is, there were no conditional relationships between geographic overlaps and floral trait differentiation (Fig. 3).

**Table 1.** The proportions of taxa pairs showing geographic isolation and floral trait isolation in the three major clades (mainland Japan, Ryukyu-Taiwan, and mainland China). The values within parentheses show the number of pairs showing overlap/differentiation and number of all pairs.

| Attribute | Clade |  |  |
| --- | --- | --- | --- |
|  | Mainland Japan | Ryukyu-Taiwan | Mainland China |
| Geographic isolation | 90.36% (704/780) | 88.60% (285/325) | 69.85% (95/136) |
| Flowering time isolation | 52.44% (409/780) | 21.37% (75/351) | 28.76% (44/153) |
| Calyx width length isolation | 36.28% (283/780) | 56.13% (197/351) | 68.63% (105/153) |

**Fig. 3.**
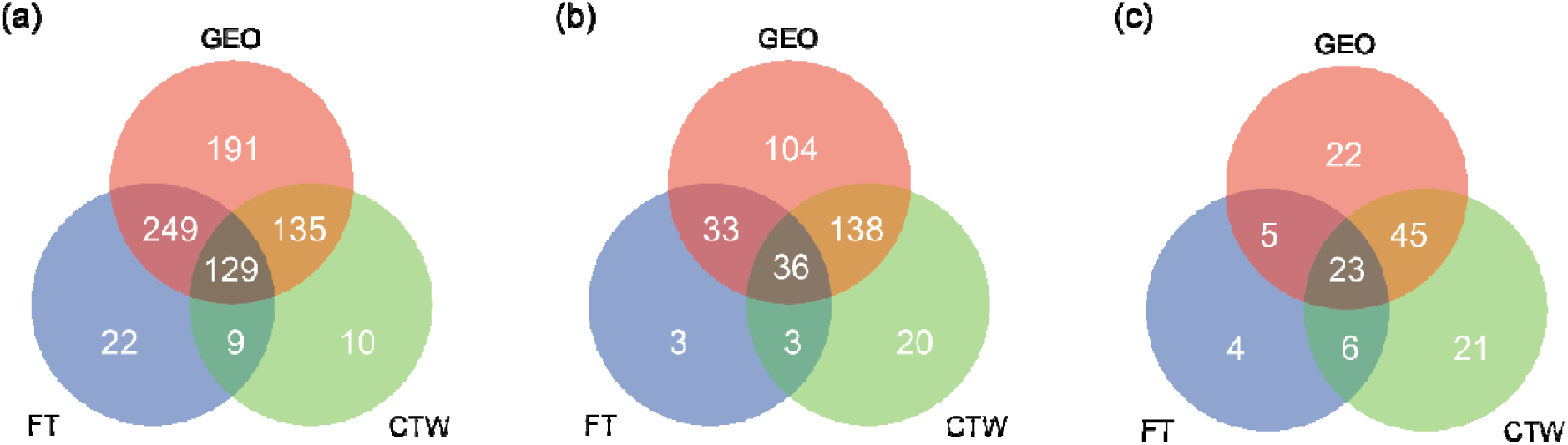
Venn diagrams showing the number of taxa pairs with isolation of distribution range (GEO; red), and flowering time (FT; blue) and calyx tube width (CTW; green) within mainland Japan clade (a), Ryukyu-Taiwan clade (b), and mainland China clade (c).

### Sister and non-sister taxa pair comparisons

Eleven sister pairs out of 24 showed geographic overlaps to various degrees (17-100%), and seven sister-pairs showed more than 80% range overlap (Table 2 & Fig. S7d). We found 14 sister pairs showing the isolation of either flowering time or calyx tube width. Within five pairs out of the 14 pairs, both traits were differentiated. In addition, six sister-pairs showed floral trait isolation with geographic overlap (for example, *Asarum blumei* vs. *Asarum nipponicum, Asarum kinoshitae* vs. *Asarum rigescens* var. *rigescens, Asarum gelasinum* vs. *Asarum monodoriflorum, Asarum celsum* vs. *Asarum gusk, Asarum yaeyamense* vs. *Asarum taipingshanianum, A. wulingense* vs. *Asarum insigne*, and *Asarum splendens* vs. *Asarum inflatum*). Our GLMM analysis showed that, of the three attributes (geographic isolation, flowering time, and calyx tube width), only geographic isolation showed higher values between non-sister taxa pairs than sister pairs (sister pairs; estimated median = 0.523, 95%CI = 0.255 – 0.7659, non-sister pairs; estimated median = 0.918, 95%CI = 0.873 – 0.954: Table S5). The other attributes showed no differentiation between sister and non-sister taxa pairs (Fig. 4). The datasets containing sister-pairs with >99% supports showed the same tendencies (Fig. S8).

**Table 2.**
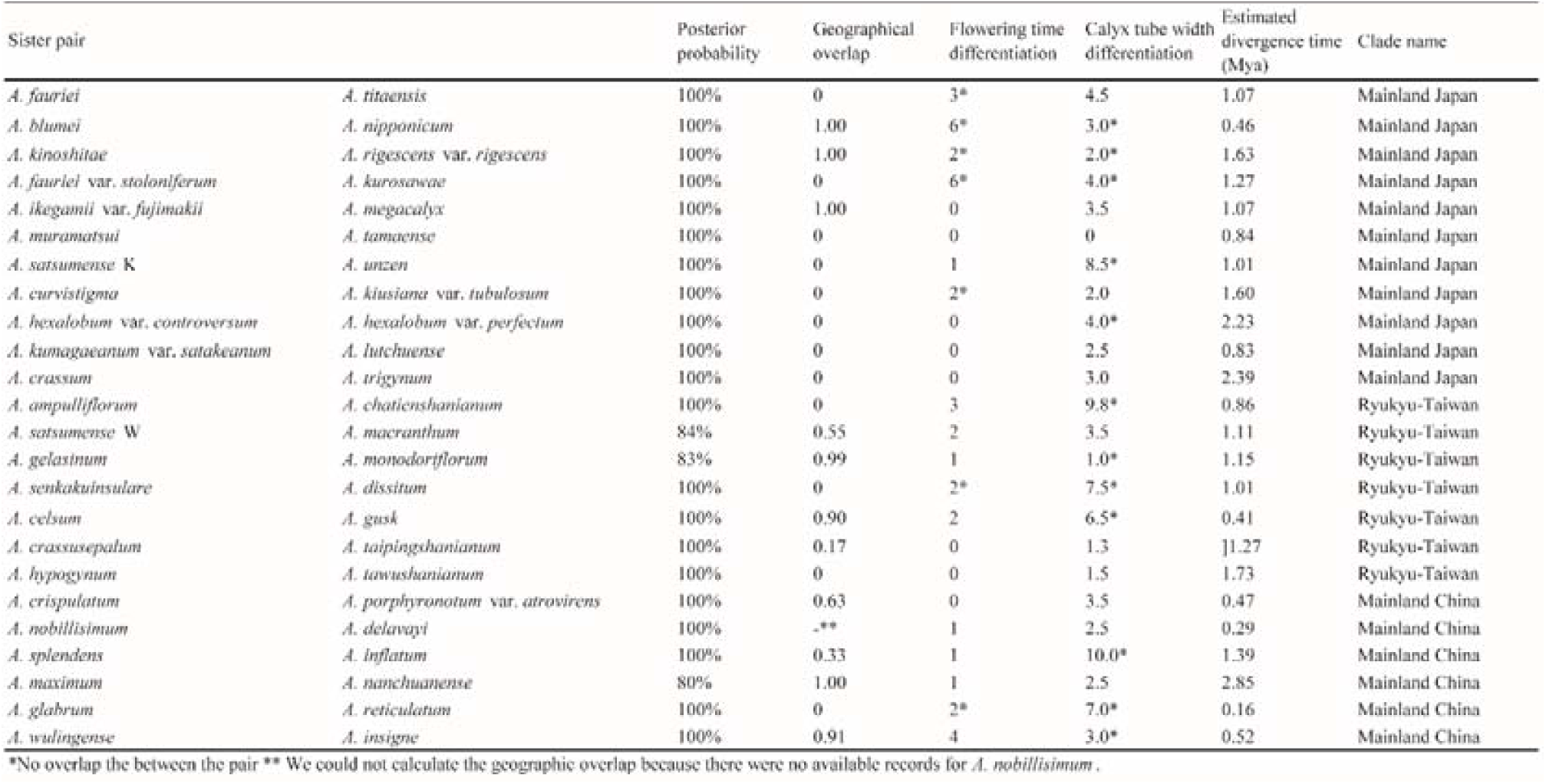
Geographic overlaps and morphological differences in the 24 sister-taxa pairs with more than 80% posterior support. For each sister-taxa pair, columns indicate the posterior probability that the two taxa are sister, proportion of range overlap, flowering time differentiation (month), calyx tube width differentiation (mm), estimated divergence time, and the clade name that includes two taxa.

**Fig. 4.**
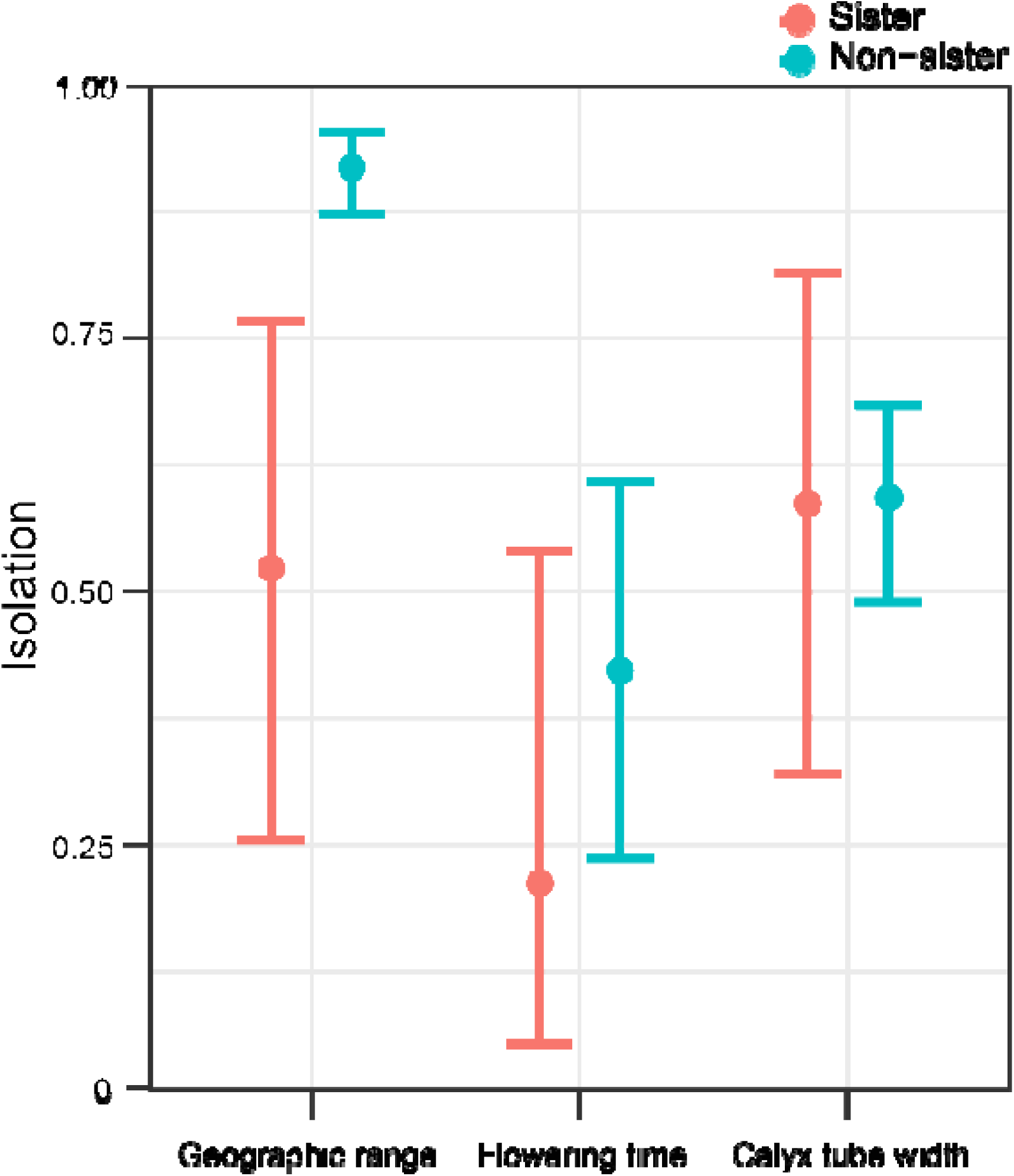
The results of sister and non-sister pairs comparisons of geographic, flowering time, and calyx tube width isolations estimated from the GLMM analysis. The mean values and 95% CIs of sister pairs and non-sister pairs are shown in red and blue, respectively. The included sister-pairs had posterior probabilities >80%.

## Discussion

### Phylogeographic history of *Heterotropa* in the SJFR

Our phylogenetic analysis using ddRAD-seq data with comprehensive sampling revealed the phylogeographic history of *Heterotropa* in the SJFR. The section originated in mainland China (Fig. 1), and the divergence time between mainland China and insular clades estimated from the chloroplast phylogenetic analysis was 9.16 Mya (95%HPD 4.66 – 13.55 Mya), corresponding to the late Miocene (Fig. S2). These results were concordant with the estimation of our previous study using the average substitution rate of the ITS region (9.3 Mya; Takahashi & Setoguchi, 2018). During the late Miocene, regressions and transgressions of the East China Sea (Haq, Hardenbol, & Vail, 1987) caused land-bridge formation that allowed the migration of temperate plants between the mainland China and insular systems (Cao, Comes, Sakaguchi, Chen, & Qiu, 2016; Yang et al., 2017). Furthermore, our study revealed that besides the insular clade, the mainland China clade was also composed of subclades, which corresponded to taxa geographic distribution (Fig. 1), and all subclades diverged during the late Miocene or the Pliocene (Fig. S3). During this period, the establishment of a monsoon climate caused by the uplift of the Himalayas and the Tibetan Plateau led to vegetational shifts in the SJFR, and frequent glacial-regressions and inter-/after-glacial transgressions (Kimura, 1996; Zheng, Powell, Rea, Wang, & Wang, 2004). We considered that geographic and climatic events would have allowed colonisation of insular systems and formation of regional lineages of *Heterotropa*, as shown in other studies (R.-S. Lu et al., 2019; Mitsui et al., 2008).

As an exception, two Chinese species were not included in the mainland China clade: *A. ichangense* and *A. forbesii* were included in or sister to the insular clade, respectively. The phylogenetic placement of the two species is consistent with a previous phylogenetic study (Okuyama et al., in press) and previous chromosomal studies that showed they have the same chromosome numbers (2n = 24) as the insular taxa, which is different from the other mainland China species (2n = 26) (Kelly, 1997; Sugawara & Ogisu, 1992). Our study using exclusive sampling of Chinese taxa demonstrated that only one back dispersal event from the insular systems to mainland China would have occurred. In addition, colonisation to other regions after formation of the regional lineages was observed only in several taxa (e.g., *A. minamitanianum* from mainland Japan to Kyushu island, and *A. lutchuense* from Kyushu island to Amami islands). This indicated that the regional lineages would have remained separated even during the Pleistocene climatic oscillations. One of the isolation mechanisms would be glacial isolation. In the SJFR, the existence of multiple refugia of temperate plants is implied by phylogeographic studies (Qiu et al., 2011). In mainland China, in addition to southern areas (< 30°N), the region around the Sichuan basin is inferred refugia and isolated due to its complex topography, including high mountains and the Yangtze River (Qiu, Guan, Fu, & Comes, 2009; Wang et al., 2015). In mainland Japan, during the glacial periods, most parts were covered by mixed (boreal and cool temperate) forests or boreal forests (Harrison et al., 2001), and warm temperate plants would have been forced to retreat southward and survive separately in narrow glacial refugia on the southern coasts of Kyushu, Shikoku, and Honshu islands (Aoki et al., 2019; Liu, Takeichi, Kamiya, & Harada, 2013). Another isolating factor would be the seaways barriers. In the Ryukyu islands, two deep-water passages (Tokara Tectonic Strait and Kerama gap, currently > 1000m in depth), were formed during the Pliocene (Kimura, 1996). These deep-water passages act as isolation barriers for plant expansion (Nakamura, Suwa, Denda, & Yokota, 2009). We considered that these glacial and geographic isolations would prevent the colonisation of most *Heterotropa* taxa.

### Diversity anomaly and its driving forces

There were no conditional relationships between geographic isolation and floral trait isolation (Fig. 3), indicating that most pairs showing floral trait isolation were distributed allopatrically. The results of trait evolutionary analyses indicated that repeated trait evolutions would have occurred after the formation of regional lineages during the Pleistocene period and that higher evolutionary rates of flowering time in mainland Japan clade and of calyx tube width in the Ryukyu-Taiwan clade were detected (Fig. 2). During the Pleistocene period, the repeated exposure and submergence of the land-bridges would have led to significant population declines and isolations of temperate plants in insular systems of the SJFR, while the populations in mainland China would have been relatively stable (Qiu et al., 2011). Considering the current smaller distribution ranges (Fig. S6) and a higher proportion of pairs showing geographic isolation (Table 1), insular *Heterotropa* taxa would have experienced repeated range fragmentations and contractions during the Pleistocene. The random genetic drifts in the range contractions could be one of the mechanisms of trait evolution in plants (Lande, 2000), and a simulation study also implied that a small population size would promote floral evolution, including flowering time without selective agents (Devaux & Lande, 2008). The neutral drift caused by the range fragmentations and contractions due to climatic oscillations would have played a role in generating the floral diversity of insular *Heterotropa*. In addition, we propose that pollinator adaptation would have also occurred through range fragmentations and contractions. The pollinator-mediated diversification of *Heterotropa* has been proposed in a previous study (Sinn et al., 2015). Empirical studies have implied that the various Diptera species are pollinators of insular *Heterotropa* taxa with specialisation (e.g., fungus gnus in *A. tamaense* [Sugawara, 1988], sciarid flies in *A. costatum* [Kakishima and Okuyama, 2018], and Calliphoridae flies in *Asarum fudsionoi* [Maeda, 2013]). These Diptera species differed in seasonal occurrence and would be attracted by different cues (Funamoto, 2019). Although we cannot deny the possibility that the pollinator difference results from the difference in the regional fauna, we consider that the floral traits of *Heterotropa* would be related to the attraction of Diptera species. Previous studies implied that geographic isolations often preceded ecological divergence, especially in low mobility organisms (Boucher et al., 2016; Heinicke, Jackman, & Bauer, 2017). In the SJFR, it was reported that the morphological heterogeneity was facilitated by geographic isolations due to Pleistocene climatic oscillations (Gao, Zhang, Gao, & Zhu, 2015), and the diversification within the insular systems was due to genetic drifts caused by range fragmentations (Yoichi et al., 2017). Thus, we considered that both selective and neutral agents facilitated by the range fragmentations during the Pleistocene period would have led to the parallel evolution of floral traits within the regional lineages and the formation of diversity anomaly of *Heterotropa* in insular systems.

### Biotic and abiotic drivers of sister taxa divergence

The results of sister and non-sister taxa comparison showed that the number of geographically overlapping pairs was higher in sister-taxa pairs (Fig. 4), indicating that, besides allopatric speciation, speciation on a small spatial scale would have also occurred in *Heterotropa* taxa. Geographical overlap between close relatives requires some kind of reproductive isolation to maintain the species boundary (Weber & Strauss, 2016). Six geographically overlapping sister taxa pairs showed floral trait isolation (Table 2). In *Heterotropa*, most taxa inhabit almost the same environments (understory of warm temperate forests) and the floral difference and/or geographic isolation would act as reproductive barriers rather than habitat differences. This insight was corroborated in a study of nine closely related *Heterotropa* taxa in the Amami islands, which are distributed in sympatry and/or close parapatry and morphologically different in floral traits (Matsuda, Maeda, Nagasawa, & Setoguchi, 2017). Thus, we considered that the flowering time and calyx tube width would have acted as one of the possible reproductive barriers in the six sister taxa pairs, and there are possibilities that the speciation on a small spatial scale triggered by trait differentiations may have also occurred in *Heterotropa*.

### Conclusion

Our results implied that the diversity anomaly in *Heterotropa* in the SFJR would have been formed by multiple drivers, including the geographic isolation and complemental floral trait evolution with different temporal scales. Phylogenetic analysis showed that in *Heterotropa*, the regional lineages would have been formed by geographic events during the Miocene, and geographic and refugial isolation would have prevented most chances of colonisation of the regional lineages. Traits evolution analysis implied that the parallel floral trait divergence associated with reproductive isolation would have occurred during the Pleistocene and that repeated range fragmentations in insular systems would have promoted the evolution of floral traits, especially in insular systems. Furthermore, the speciation on a small spatial scale with floral trait divergence was also implied in at least six sister taxa-pairs. Hence, the diversification of insular *Heterotropa* would have been triggered by the geographic and climatic events during the Miocene, and subsequent repeated floral trait evolution with and without geographic isolations in the regional lineages during the Pleistocene. Our study demonstrated the importance of multidimensional studies to understand the diversification process of temperate plants in the SJFR, where geographic isolation has been considered to play a major role in the diversification.

## Acknowledgements

We thank S. Gale, Y. Inoue, S. Liao, K. Maeda, J. Nagasawa, S. Nemoto, T. Teramine, Y. Wakita, M. Yamamoto, and S. Zhou for their help with sampling and taxonomic determination. We are also grateful to J. R. P. Worth, R. Imai, and M. Yamasaki for their valuable comments on the genetic and statistical analyses and manuscript writing. This work was supported by Grants-in-Aid for Scientific Research from the Japanese Society for the Promotion of Science (Nos. 24247013, 26304013 and 18J22919), the Environment Research and Technology Development Fund (grant no. 4-1702 and 4-1902), and the Environmental Research and Technology Development Fund of the Ministry of the Environment SICORP Program of the Japan Science and Technology Agency (“Spatial-temporal dimensions and underlying mechanisms of lineage diversification and patterns of genetic variation of keystone plant taxa in warm-temperate forests of Sino-Japanese Floristic Region”) (grant no. 4-1403). We would like to thank Editage (www.editage.com) for English language editing.

## Data accessibility

The obtained reads for chloroplast genome construction and ddRAD-seq analysis are available in NCBI (GenBank BioProject no. PRJDB9302 and PRJDB8943) The alignment sequences, morphological data, and distribution records were deposited in the Dryad^®^ digital repository under doi: 10.5061/dryad.xwdbrv1b4.

## Author contributions

D. T., S. S., Y. Q., Y. I., and H. S. conceptualised and designed the study. Sample collection was performed by D. T., S. S., Y. F., Y. Q., Y. I., P. L., R. L., C. L., S. C., Y. L., Y. C., and H. S. The molecular experiments were conducted by D. T., S. S., L. K., and A. N., Data were generated, analysed, and visualised by D. T. and S. S. Manuscript writing was led by D. T., with contributions from all authors.

## Conflict of interest

The authors declare that they have no conflicts of interest.

## Supplemental Figures and Tables

Fig. S1. Part of the floral diversity of *Heterotropa* taxa. The font and the side views of the flowers of *Asarum takaoi* (a), *A. savatieri* var. *iseanum* (b), *A. nipponicum* (c), *A. hexalobum* var. *perfectum* (d), *A. rigescens* (e), *A. costatum* (f), *A. trigynum* (g), *A. sakawanum* var. *stellatum* (h), *A. satsumense* K (i), *A. dissitum* (j), *A. pellucidum* (k), *A. gusk* (l), *A. senkakuinsulare* (m), *A. yaeyamense* (n), *A. tokarense* (o), *A. villisepalum* (p), *A. hypogynum* (q), *A. chatienshanianum* (r), *A. macranthum* (s), *A. forbesii* (t), *A. ichangense* (u), *A. delavayi* (v), *A. petelotii* (w), *A. inflatum* (x), and *A. maximum* (y). The taxa were ordered according to their distributions: mainland Japan (a-h), the Ryukyu Islands (i-o), Taiwan (p-s), and mainland China (t-y). The colours of the alphabet indicate the flowering time of the taxa (red; autumn, blue; winter, and green; spring).

Fig. S2. Phylogenetic tree based on 59 CDS regions (26,786bp) of the chloroplast genomes of the Magnoliids and Chloranthales species. The red circle indicates the calibration point (169 – 180 Mya) followed by Zeng et al. (2014). The bars indicated 95% highest posterior density (HPD) intervals of estimated divergence times of nodes. The posterior probabilities of all the blanches were > 0.999.

Fig. S3. A majority-rule consensus tree inferred from Bayesian analysis of ddRAD-seq data (50% genotyped data) showing the divergence times of major nodes and 95% highest posterior density (HPD) intervals.

Fig. S4. Molecular phylogenetic tree of *Heterotropa* taxa estimated from Bayesian analysis using 75% genotyped (a) and 90% genotyped (b) matrices. Values above or below branches indicate the posterior probabilities of branches.

Fig. S5. Histograms of supported number of macroevolutionary rate regimes for each trait evolution; (a) flowering time and (b) calyx tube width. The number of regimes = 1 indicated there were no shifts in trait evolution rate throughout the tree.

Fig. S6. Histograms of distribution area of taxa within mainland Japan clade (a), Ryukyu-Taiwan clade (b), and mainland China clade (c).

Fig. S7. Histograms of proportion of geographic range overlap between taxa pairs within mainland Japan clade (a), Ryukyu-Taiwan clade (b), mainland China clade (c), and sister-taxa pairs (d).

Fig. S8. The results of sister and non-sister taxa pairs comparisons of the three attributes (geographical, flowering time, and calyx width isolations), including sister taxa pairs supported by more than 99% posterior probabilities. The mean values and 95% CIs of sister pairs and non-sister pairs are shown in red and blue, respectively.

Table S1. Sample list used in ddRAD-seq analysis. The columns indicate the distribution region used in S-DIVA, the median value of flowering time (month), median value of calyx tube width (mm), distribution area calculated from the polygon formed by the distribution record(s), number of records used in the geographic analysis, the number of cleaned RAD-seq reads, and the number of loci in the 50% matrices.

Table S2. Information of newly obtained reads and chloroplast genomes of *Asarum* spp. Table S3. The sample lists used in phylogenetic analysis of chloroplast genome.

Table S4. The values of κ and λ Pagel’s statistics for flowering time and calyx tube width calculated from the phytools package.

Table S5. The estimated median values and 95% CIs of each attribute (geographic, flowering time, and calyx tube width isolations) in sister pairs with posterior probabilities of nodes higher than 80% and 99%, and non-sister pairs.

## Notes

### Competing Interest Statement

The authors have declared no competing interest.

