## Supplementary figures and images for "Geographic and subsequent biotic isolations led to a diversity anomaly of *Heterotropa* (Aristolochiaceae) in insular versus continental regions of the Sino-Japanese Floristic Region"

### Supplementary Figures

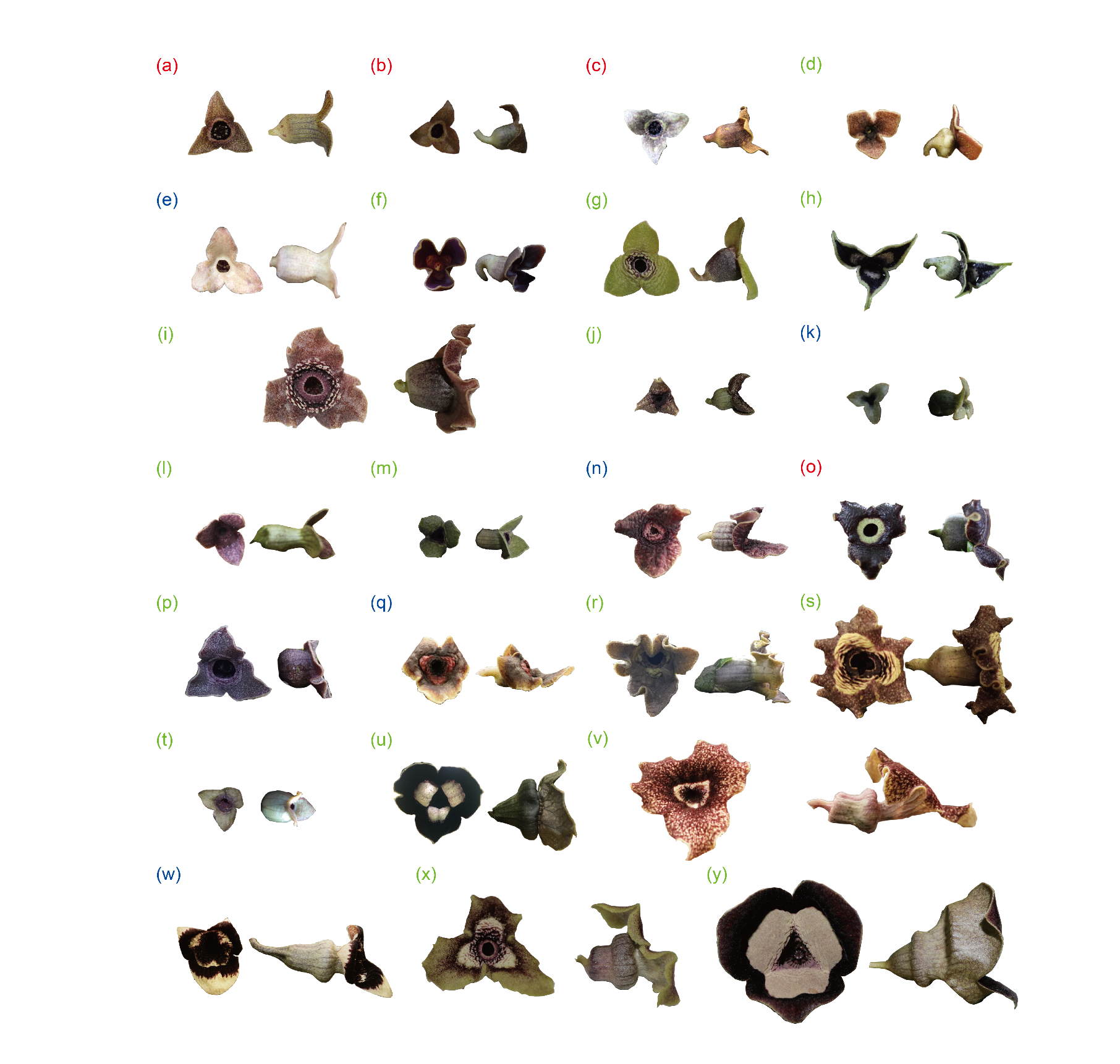
Fig. S1

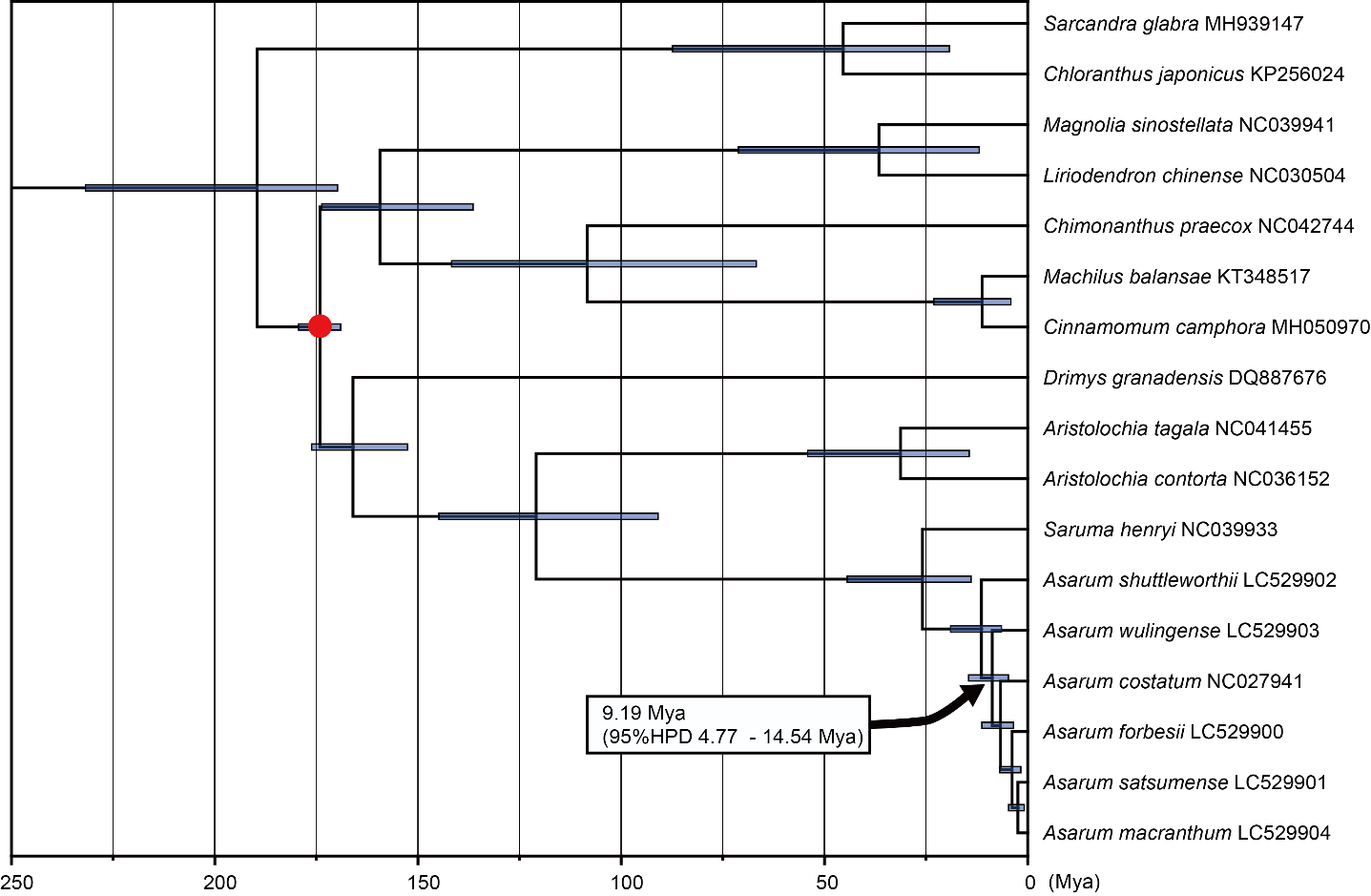
Fig. S2

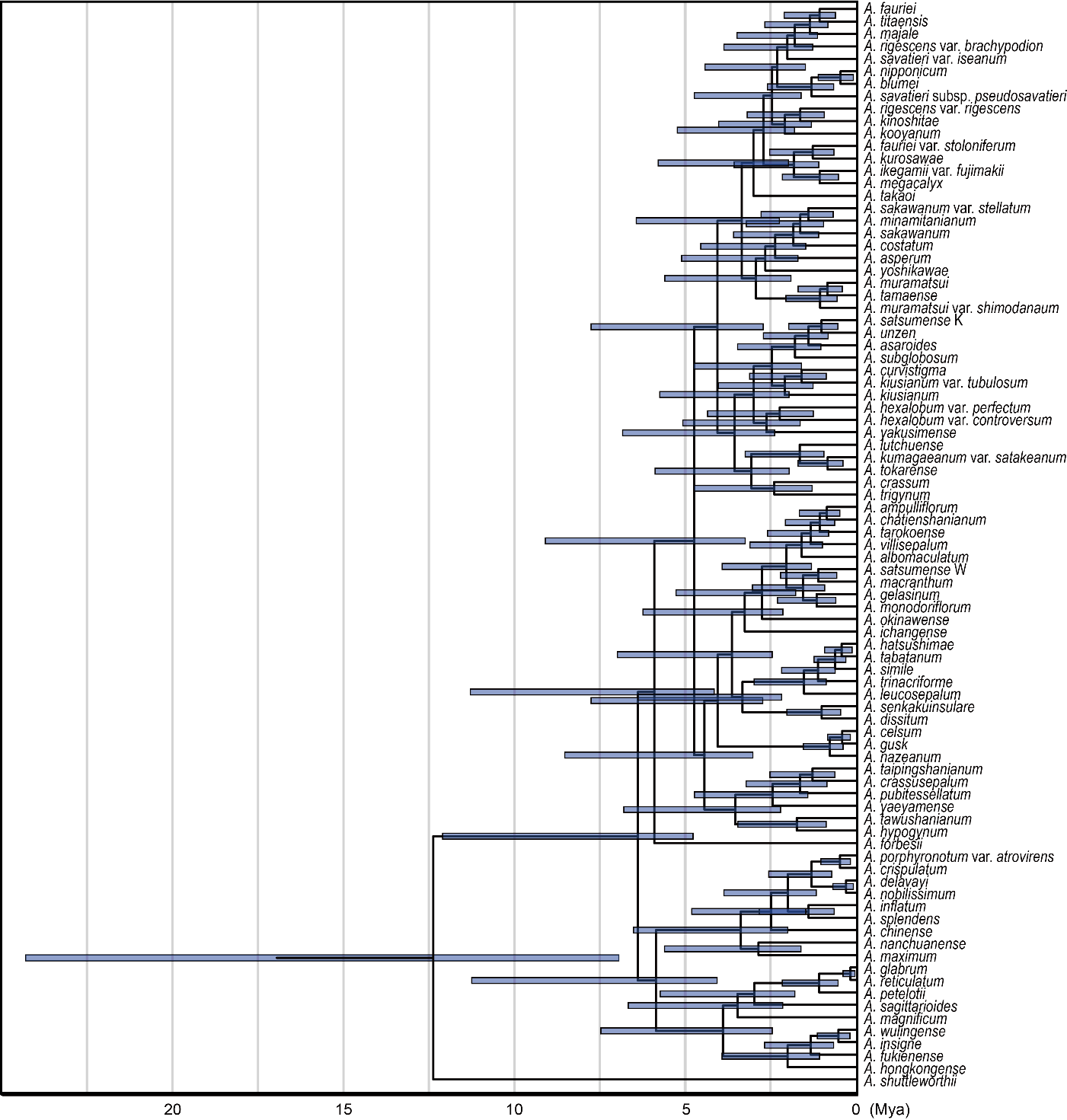
Fig. S3

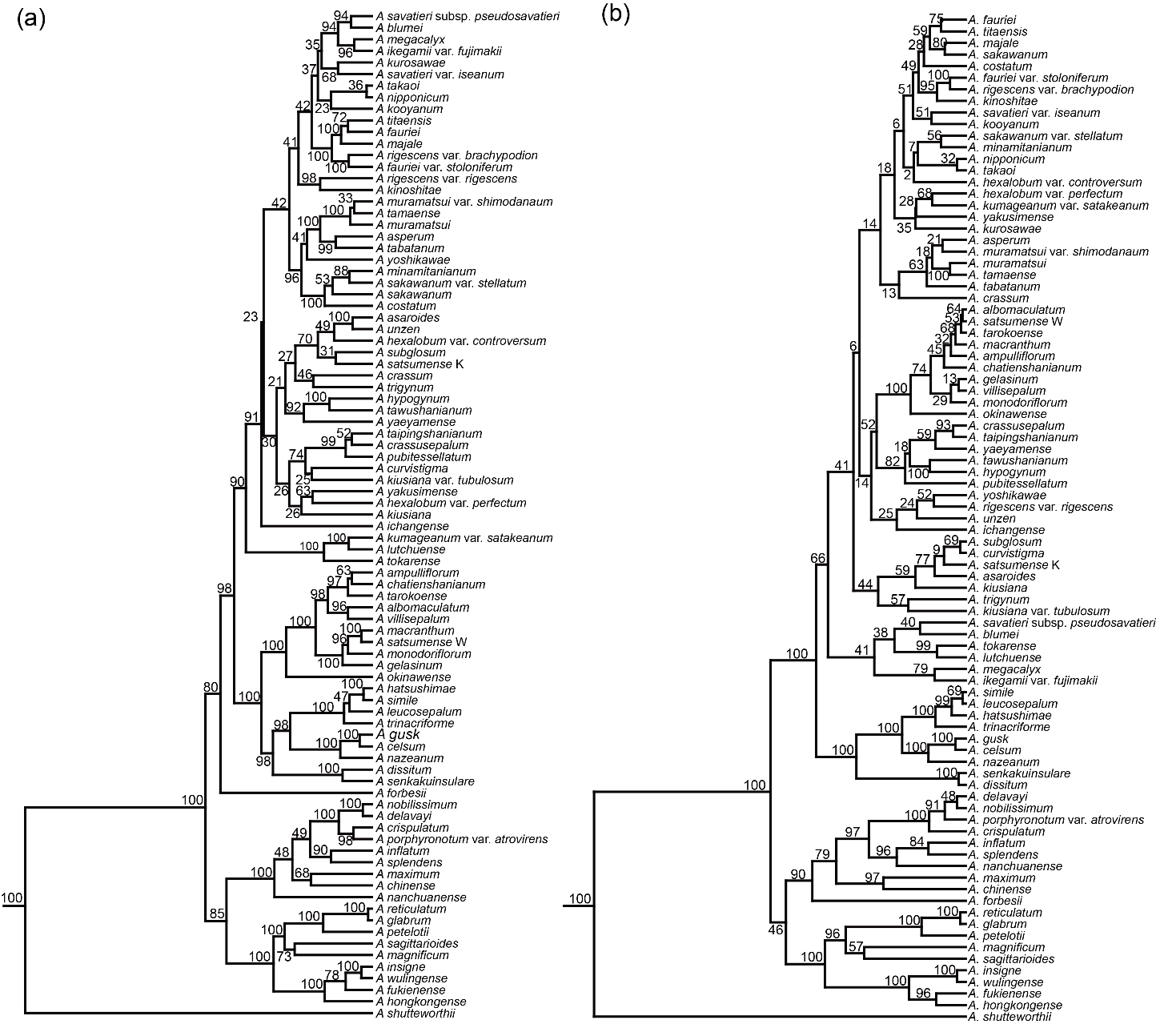
Fig. S4

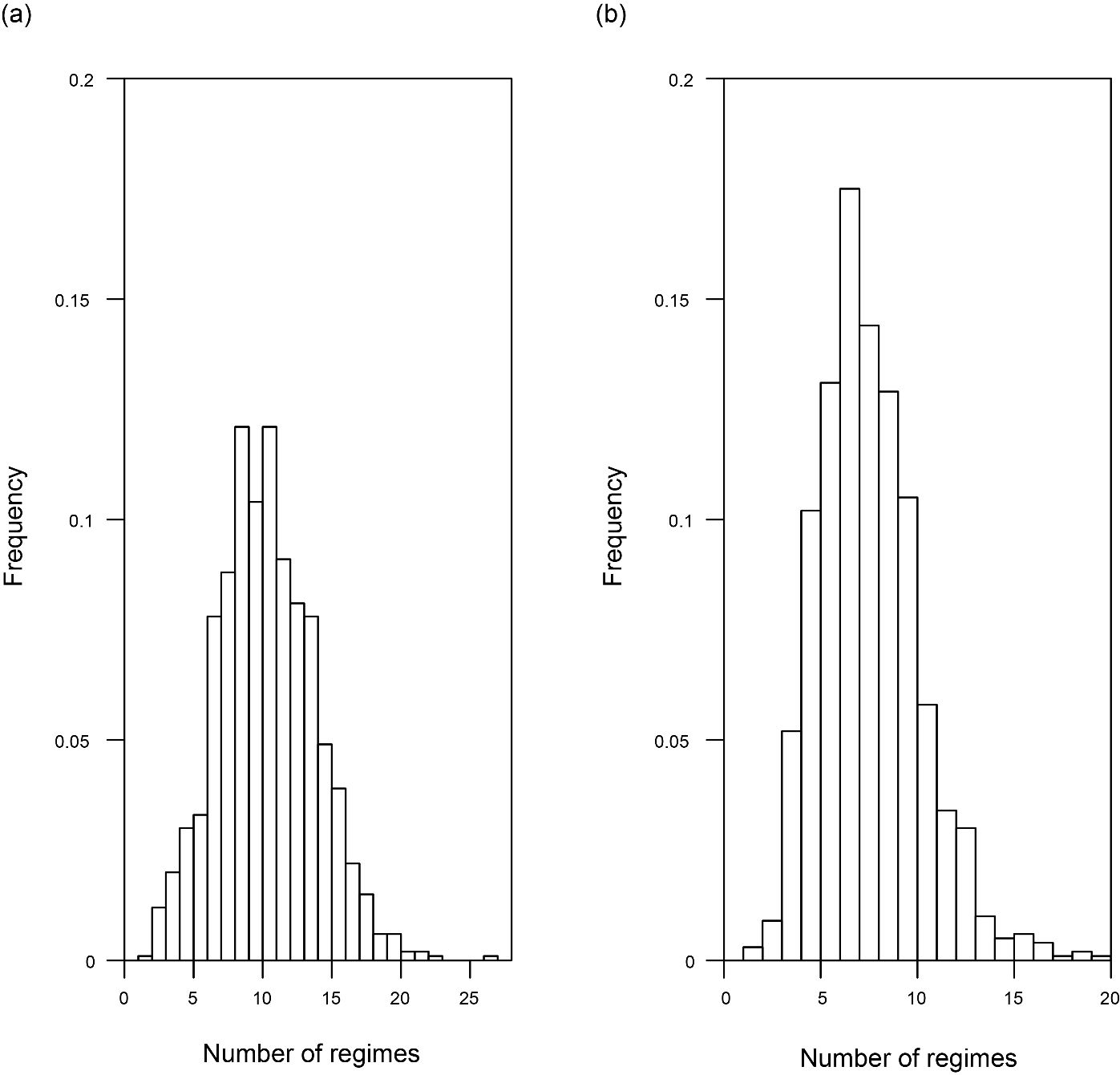
Fig. S5

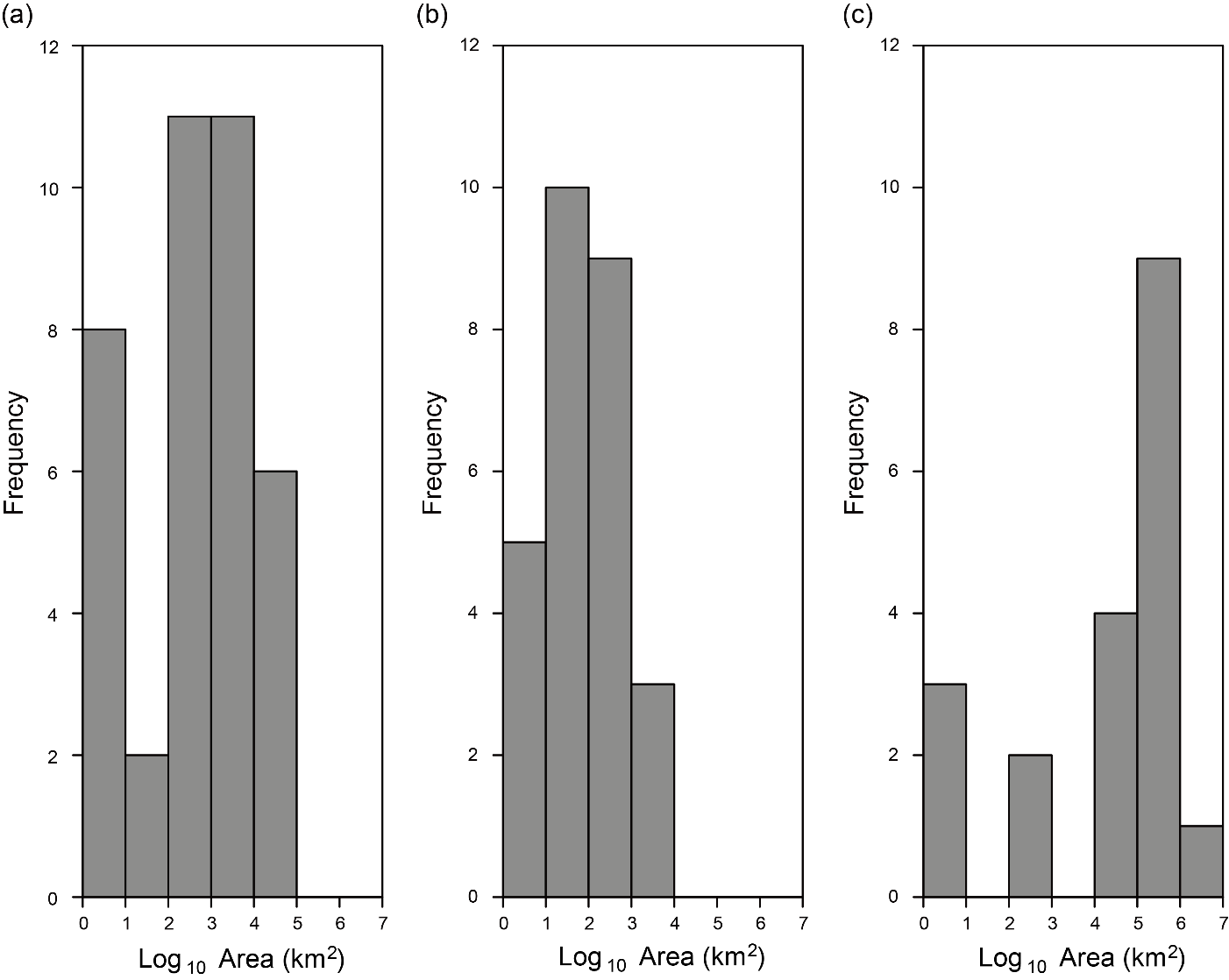
Fig. S6

Fig. S7

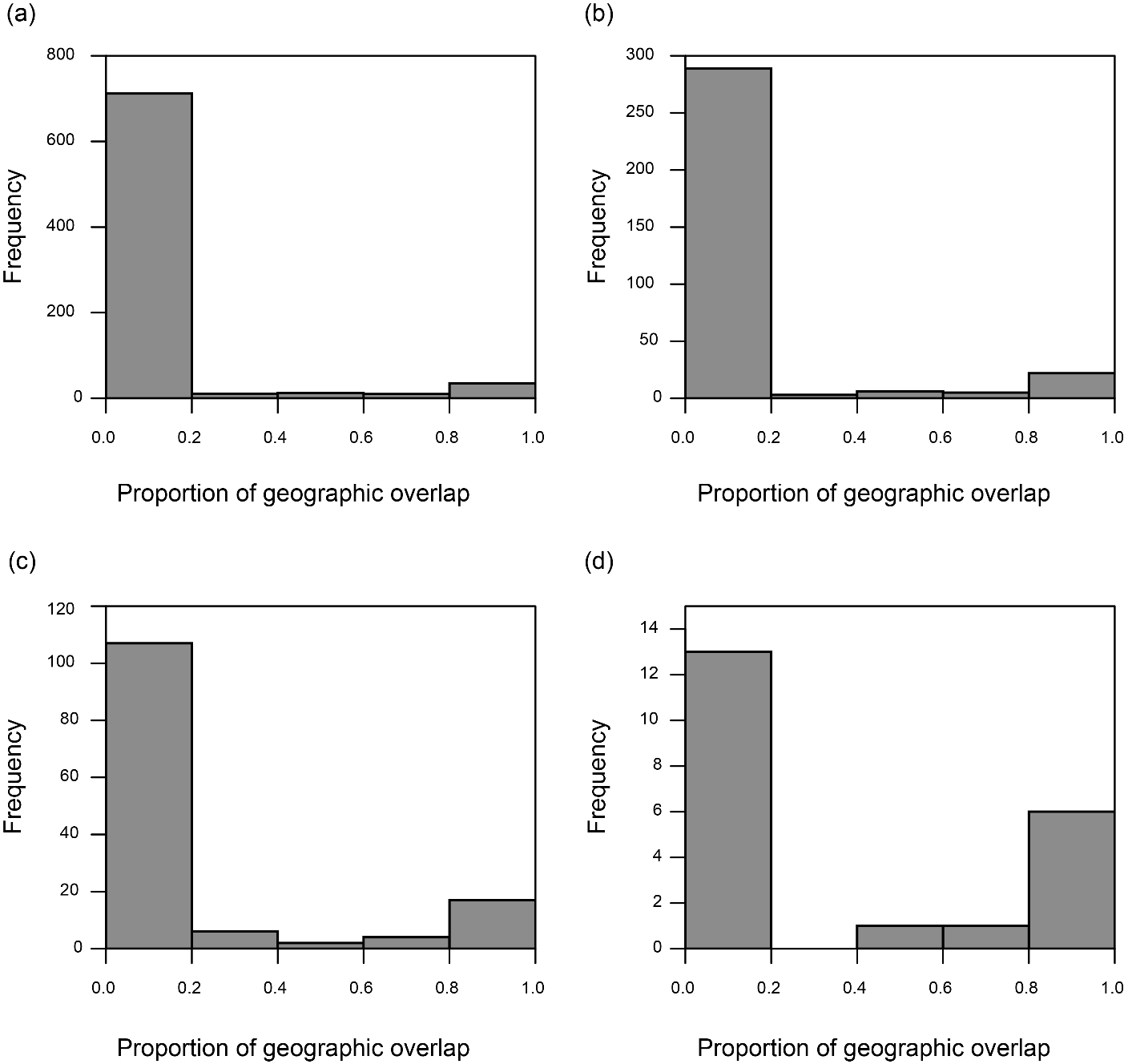

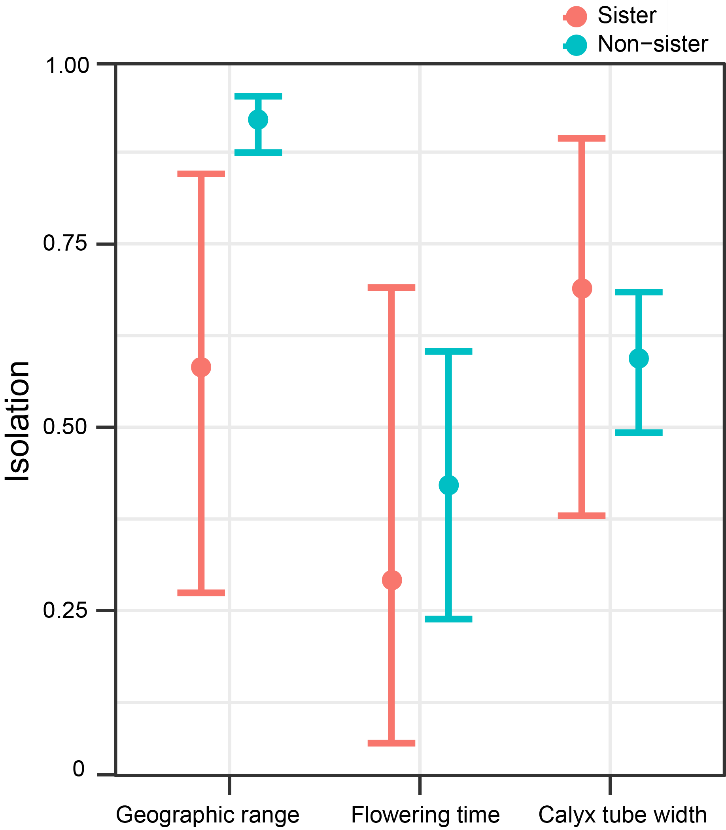
Fig. S8
