## Appendix 1 for "Geographic and subsequent biotic isolations led to a diversity anomaly of *Heterotropa* (Aristolochiaceae) in insular versus continental regions of the Sino-Japanese Floristic Region"

***Chloroplast genome construction and divergence time estimation***

To obtain the chloroplast genomes, we sequenced four *Heterotropa* species (*Asarum satsumense* K, *Asarum macranthum*, *Asarum wulingense*, and *Asarum forbesii*) and one *Hexastylis* species (*Asarum shuttleworthii*). We used three sequencing methods.

To *A. macranthum* and *A. wulingense*, we used the chloroplast enrichment method following Sakaguchi *et al.* (2017). To construct barcoded DNA fragment libraries, the Ion Xpress Plus Fragment library Kit (Thermo Fisher Scientific, Waltham, Massachusetts, USA) was used to process the purified DNA of *A. macranthum* and *A. wulingense*. The barcoded libraries were mixed with Ion Sphere Particle for emulsion PCR using Ion One Touch 2 system (Thermo Fisher Scientific) with Ion PGM Hi-Q OT2 Kit (Thermo Fisher Scientific). From the product of emulsion PCR, the positive particles with amplified DNA were isolated and purified by Ion OneTouch ES (Thermo Fisher Scientific) and loaded onto an Ion 318 chip (Thermo Fisher Scientific). Sequencing was performed using an Ion PGM sequencer (Thermo Fisher Scientific). Extracted DNA from *A. satsumense* K and *A. shuttleworthii* were fragmented using the Takara DNA Fragmentation Kit (Takara Bio, Ohtsu, Shiga, Japan). The library preparation was conducted using the SMARTer ThruPLEX DNA-Seq Kit (Takara Bio, USA). Barcoded libraries were sequenced with paired-end 150 bp reads on Illumina Hiseq-X (Illumina, San Diego, California, USA; sequencing was performed by Macrogen Japan, Kyoto, Japan). To *A. forbesii*, NEBNext® DNA Library Prep Kit (New England BioLabs, Ipswich, MA, USA) was used to prepare library. Nova-seq 6000 (Illumina) was used to sequence the prepared library. Library preparation and sequencing were performed by Chemical Dojin (Kumamoto, Japan).

All obtained reads were trimmed using Trimomomatic v. 0.32 software (Bolger *et al.*, 2014) using the following commands: HEADCRAP:10, LEADING:20, TRAILING:20, SLIDINGWINDOW:4:20, AVGQUAL:20, and MINLEN:50. Because the amount of obtained reads of *A. forbesii* was too large (> 80 Gb), we reduced the data to 500,000 reads (approximately 7.5 Gb). The cleaned reads were mapped using MITObim v.1.8 (Hahn *et al.*, 2013) to the chloroplast genome of *Asarum costatum* (AP018513; Takahashi *et al.*, 2018) with minimum depth 4X. The obtained reads and chloroplast genomes were deposited in DDBJ (BioProject ID, PRJDB9302, Table S2).

To construct chloroplast genome phylogeny and estimate divergence time, in addition to newly obtained five sequences, we used the chloroplast genome sequences of ten Magnoliid species, including one *Heterotropa* species (*Asarum costatum*), and two Chloranthales species. The species information is shown in Table S3. As our data set included highly divergent species, to construct the chloroplast phylogeny, we used only CDS regions shared by more than 17 species out of 18 species. The CDS regions of chloroplast genomes and assemblies were identified using GeSeq with protein search identity value 85 (Tillich *et al.*, 2017). Sequence data were manually edited and aligned using BioEdit v.7.0.5.3 (Hall, 1999). In total, 59 regions (26,786 bp) were used for the phylogenetic analysis. The phylogenetic tree construction and estimation of divergence time was conducted by using BEAST v.10.0.4 (Drummond & Rambaut, 2007) applying the GTR+I+G model inferred by JmodelTest v.1.4.7 (Posada, 2008). To estimate the divergence time of the crown age of the *Heterotropa* clade, the crown of Magnoliids was constrained using a uniform distribution with a lower bound of 169 Mya and upper bound of 180 Mya according to the study of angiosperm phylogeny using fossil calibrations (Zeng *et al.*, 2014). The Markov Chain Monte Carlo method was performed using four independent runs with four chains of 50,000,000 generations each, saving one tree every 1000 generations. The first 10,000,000 generations were discarded as burn-in, as evaluated by TRACER v.1.5 (Rambaut & Drummond, 2013). The obtained tree was displayed using FigTree v.1.4 (Rambaut, 2009).

***Reference lists***

**Bolger AM, Lohse M, Usadel B. 2014.** Trimmomatic: a flexible trimmer for Illumina sequence data. *Bioinformatics* **30**(15): 2114-2120.

**Drummond AJ, Rambaut A. 2007.** BEAST: Bayesian evolutionary analysis by sampling trees. *Bmc Evolutionary Biology* **7**: 8.

**Hahn C, Bachmann L, Chevreux B. 2013.** Reconstructing mitochondrial genomes directly from genomic next-generation sequencing reads-a baiting and iterative mapping approach. *Nucleic Acids Research* **41**(13).

**Hall TA 1999**. BioEdit: a user-friendly biological sequence alignment editor and analysis program for Windows 95/98/NT. *Nucleic acids symposium series*. 95-98.

**Posada D. 2008.** jModelTest: Phylogenetic model averaging. *Molecular Biology and Evolution* **25**(7): 1253-1256.

**Rambaut A 2009**. FigTree v1. 4: Tree figure drawing tool.

**Rambaut A, Drummond AJ 2013**. Tracer v1. 5 Available from <http://beast>. bio. ed. ac. uk/Tracer: Accessed.

**Sakaguchi S, Ueno S, Tsumura Y, Setoguchi H, Ito M, Hattori C, Nozoe S, Takahashi D, Nakamasu R, Sakagami T, et al. 2017.** Application of simplified method of chloroplast enrichment to small amounts of tissues for chloroplast genome sequencing. *Applications in Plant Sciences* **5**(5).

**Takahashi D, Sakaguchi S, Isagi Y, Setoguchi H. 2018.** Comparative chloroplast genomics of series *Sakawanum* in genus *Asarum* (Aristolochiaceae) to develop single nucleotide polymorphisms (SNPs) and simple sequence repeat (SSR) markers. *Journal of Forest Research* **23**(6): 387-392.

**Tillich M, Lehwark P, Pellizzer T, Ulbricht-Jones ES, Fischer A, Bock R, Greiner S. 2017.** GeSeq - versatile and accurate annotation of organelle genomes. *Nucleic Acids Research* **45**(W1): W6-W11.

**Zeng LP, Zhang Q, Sun RR, Kong HZ, Zhang N, Ma H. 2014.** Resolution of deep angiosperm phylogeny using conserved nuclear genes and estimates of early divergence times. *Nature Communications* **5**.
